# Arabidopsis *XTH4* and *XTH9* contribute to wood cell expansion and secondary wall formation

**DOI:** 10.1101/2019.12.16.877779

**Authors:** Sunita Kushwah, Alicja Banasiak, Nobuyuki Nishikubo, Marta Derba-Maceluch, Mateusz Majda, Satoshi Endo, Vikash Kumar, Leonardo Gomez, Andras Gorzsas, Simon McQueen-Mason, Janet Braam, Björn Sundberg, Ewa J. Mellerowicz

**Author notes:** Corresponding Author Dr. Ewa J. Mellerowicz, Professor, Department of Forest Genetics and Plant Physiology, Swedish University of Agricultural Sciences (Sveriges lantbruksuniversitet), S901-83 Umeå, Sweden. Equal contribution (Sunita Kushwah, Alicja Banasiak). **AUTHOR CONTRIBUTION** SK analyzed cell wall composition, mutant growth, and gene expression, and wrote the manuscript; AB analyzed hypocotyl phenotypes and xylem cell morphology in mutants, discovered secondary wall phenotypes, and carried immunolocalization, NN isolated and purified mutants and created OE plants, MDM and MM analyzed cell walls by *in situ* Raman and *in situ* FTIR, SE created GUS reporter lines, VK analyzed *Populus* XTH genes and their expression in AspWood, LG and SMM carried out saccharification analyses, AG interpreted FTIR and Raman data and helped with spectroscopic analyses; JB analyzed XET activity, BS and EJM conceived and coordinated the project, EJM finalized the manuscript with contribution from all authors. **FUNDING INFORMATION** The financial support of the Chemical Biological Centre (KBC) and the Department of Chemistry of Umeå University for the Vibrational Spectroscopy Core Facility, Bio4Energy and TC4F support for Plant Analytical Biopolymer platform, and Vinnova (the Swedish Governmental Agency for Innovation Systems) and KAW (The Knut and Alice Wallenberg Foundation) support for plant growth facility are acknowledged. The research projects were funded by SSF (ValueTree project RBP14-0011), Formas and VR grants to EJM.

## Abstract

In dicotyledons, xyloglucan is the major hemicellulose of primary walls affecting the load-bearing framework with participation of XTH enzymes. We used loss- and gain-of function approaches to study functions of abundant cambial region expressed *XTH4* and *XTH9* in secondary growth. In secondarily thickened hypocotyls, these enzymes had positive effects on vessel element expansion and fiber intrusive growth. In addition, they stimulated secondary wall thickening, but reduced secondary xylem production. Cell wall analyses of inflorescence stems revealed changes in lignin, cellulose, and matrix sugar composition, indicating overall increase in secondary versus primary walls in the mutants, indicative of higher xylem production compared to wild type (since secondary walls were thinner). Intriguingly, the number of secondary cell wall layers was increased in *xth9* and reduced in *xth4*, whereas the double mutant *xth4x9* displayed intermediate number of layers. These changes correlated with certain Raman signals from the walls, indicating changes in lignin and cellulose. Secondary walls were affected also in the interfascicular fibers where neither *XTH4* nor *XTH9* were expressed, indicating that these effects were indirect. Transcripts involved in secondary wall biosynthesis and in cell wall integrity sensing, including *THE1* and *WAK2*, were highly induced in the mutants, indicating that deficiency in *XTH4* and *XTH9* triggers cell wall integrity signaling, which, we propose, stimulates the xylem cell production and modulates secondary wall thickening. Prominent effects of *XTH4* and *XTH9* on secondary xylem support the hypothesis that altered xyloglucan can affect wood properties both directly and *via* cell wall integrity sensing.

**SIGNIFICANCE STATEMENT:** Xyloglucan is a ubiquitous component of primary cell walls in all land plants but has not been so far reported in secondary walls. It is metabolized *in muro* by cell wall-residing enzymes - xyloglucan endotransglycosylases/hydrolases (XTHs), which are reportedly abundant in vascular tissues, but their role in these tissues is unclear. Here we report that two vascular expressed enzymes in Arabidopsis, XTH4 and XTH9 contribute to the secondary xylem cell radial expansion and intrusive elongation in secondary vascular tissues.

Unexpectedly, deficiency in their activities highly affect chemistry and ultrastructure of secondary cell walls by non-cell autonomous mechanisms, including transcriptional induction of secondary wall-related biosynthetic genes and cell wall integrity sensors. These results link xyloglucan metabolism with cell wall integrity pathways, shedding new light on previous reports about prominent effects of xyloglucan metabolism on secondary walls.

**One sentence summary:** XTH4 and XTH9 positively regulate xylem cell expansion and fiber intrusive tip growth, and their deficiency alters secondary wall formation via cell wall integrity sensing mechanisms.

## INTRODUCTION

Plant cell wall is composed of cellulose microfibrils embedded in a matrix of hemicelluloses and pectins, structural glycoproteins and, in some cell types, lignin. Xyloglucan (XG) is an abundant hemicellulose present in all lineages of plant species studied till date as well as in green algae (Popper et al., 2011). In dicotyledons, including Arabidopsis, XG constitutes approx. 20% of dry weight in primary walls, but the mature secondary walls have not been reported to contain XG (Scheller and Ulvskov, 2010).

XG backbone is made up of β-(1,4)-linked glucose substituted with α-(1,6)-linked xylose (Xyl), which can be further decorated with β-(1,2) galactose (Gal) residues with or without substitution by α-(1,2) fucose (Fuc) (reviewed by Pauly and Keegstra, 2016). Acetyl groups are usually present on Gal. XG substitutions are plant species- and organ-/tissue-/cell-type specific and can vary during cell development. In the majority of dicots, XXXG-type XGs are present where three out of four backbone residues are xylosylated, and the two last Xyl residues β-1,2 galactosylated, with the last Gal frequently α-1,2 fucosylated. In some tissues, like root hairs, galacturonic acid (GalA) can be attached to Xyl at position 2 (Peña et al., 2012).

XG coats hydrophobic microfibril surfaces (Park and Cosgrove, 2012; 2015). Earlier models suggested that XG crosslinks adjacent microfibrils (Hayashi et al., 1994). Recently, however, it was proposed to form ‘biomechanical hot-spots’ instead, which are the links between adjacent microfibrils with XG embedded in between (Park and Cosgrove, 2012; 2015). XG also covalently links with pectins (Thompson and Fry, 2000; Popper and Fry, 2005; 2008), and with ARABINOXYLAN PECTIN ARABINOGALACTAN PROTEIN1 (APAP1) (Tan et al., 2013).

XG biosynthesis starts in the Golgi apparatus as a team effort of glucan synthases encoded by *CELLULOSE SYNTHASE-LIKE C4* (*CSLC4*), three XG-xylosyltransferases (XXTs), XXT1, 2 and 5, two galactosyltransferases, MUR3 and XLT2, and a fucosyltransferase FUT1 (Pauly and Keegstra, 2016). Nascent XG chains are packaged into the secretory vesicles, and secreted into the apoplast where they are incorporated into the existing XG-cellulose network nonenzymatically and/or by the XG *endo*-transglycosylase (XET) activity. XET enzymes discovered in 1992 by three independent groups (Farkas et al. 1992; Fry et al., 1992; Nishitani and Tominaga, 1992; Okazawa et al., 1993). They are grouped in Carbohydrate-Active enZYmes (CAZY) family GH16, along with XG *endo*-hydrolases (XEHs), and thus the family has been named as XTH for **X**G *endo*-**tr**ansglycosylase/**h**ydrolase (Rose et al., 2002). The XTH family includes three major clades I/II, IIIA and IIIB, and XEH activity has been reported only in clade IIIA (Baumann et al., 2007; Eklöf and Brumer, 2010; Kaewthai et al., 2013). XETs cleave the XG backbone (donor) endolytically, and the enzyme-bound XG fragment is then transferred to the non-reducing end of another XG chain (acceptor), and the glycosidic linkage is re-formed (Eklöf and Brumer, 2010). Few XTH members are able to use other than XG or H_2_O acceptors, including mixed-link glucan (Hrmova et al., 2007; Simmons et al., 2015, Simmons and Fry, 2017), or use amorphous cellulose as acceptor and donor (Shinohara et al., 2017).

XETs are thought to regulate cell wall plasticity along with expansins and pectin-digesting enzymes but the molecular action of their activity in cell walls is not fully understood (Van Sandt et al., 2007; Park and Cosgrove, 2015). XET activity has been demonstrated to incorporate the fluorescently labeled XGOs to a fraction aligned with cellulose microfibrils, possibly by transglycosylation of the XG loose ends or cross-links, thus probably resulting in wall loosening, as well as to a cell wall fraction inaccessible to the XG degrading enzymes (Vissenberg et al., 2005a). The latter could possibly constitute the biomechanical hot-spots and XET activity would then contribute to wall strengthening. Analysis of several *xth* mutants revealed reduced cell sizes (Osato et al., 2006; Liu et al., 2007; Sasidharan et al., 2010; Ohba, et al., 2011), whereas the overexpression or exogenous application of XET proteins either stimulated (Shin et al., 2006; Ohba et al., 2011; Miedes et al., 2013) or decreased (Maris et al., 2009) cell expansion. Other studies have shown that XETs could be involved in either wall loosening or strengthening depending on the acceptor size (Takeda et al., 2002).

XETs are known to be highly expressed, both in primary (Xu et al., 1995; Antosiewicz et al., 1997; Oh et al., 1998; Dimmer et al., 2004; Vissenberg et al., 2005b; Jiménez et al., 2006; Hara et al., 2014), and in secondary (Bourquin et al., 2002; Nishikubo et al., 2007; Goulao et al., 2011) vascular tissues, but their roles in these tissues are not fully understood. Only one gene, *AtXTH27*, has been functionally characterized in vascular tissues and shown to affect tracheary element development in minor veins of rosette leaves, and proposed to mediate XG degradation in these cells (Matsui et al., 2005). In aspen, *Ptxt*XTH34 protein and XET activity were highly expressed in developing xylem fibers, coinciding with CCRC-M1 antibody (recognizing the terminal Fuc of XG) signals (Puhlmann et al., 1994), and suggesting a deposition of XG to primary wall layers (compound middle lamella) through the developing secondary cell wall layers (Bourquin et al., 2002; Nishikubo et al., 2011). In support of this hypothesis, several genes similar to XG xylosyl transferases were found expressed during secondary wall formation (Sundell et al., 2017) raising a possibility of continuous XG deposition and *XTH* function in this process. Indeed, the overexpression of *PtxtXTH34* resulted in more CCRC-M1 signals in the compound middle lamella and more cell wall-tightly bound XG at early stages of secondary xylem cell differentiation. But the later stages of xylogenesis did not show increased XG anymore and the role of such XET-induced XG deposition in xylem cells remained elusive.

To address the role of *XTH* genes during secondary xylem development, we analyzed patterns of *XTH* gene family expression in developing wood, using the AspWood database (Sundell et al., 2017). For functional analyses, we selected two Arabidopsis genes, *AtXTH4* and *AtXTH9,* similar to abundant aspen *XTH* transcripts exhibiting the most frequently observed expression pattern and tested if these two genes are involved in xylem cell expansion, or in other aspects of xylem cell differentiation. Mutant analysis revealed that *AtXTH4* and *AtXTH9* do not only regulate xylem cell expansion, but also influence several characteristics of secondary growth, including secondary xylem production and secondary wall deposition. The deficiency in these two genes was additive for some traits, suggesting their partially redundant or additive roles for those traits, while it was unique or even opposite for other traits. The upregulation of several cell wall integrity-related genes in these *xth* mutants and their non-cell autonomous effects suggest that some of them are induced by the cell wall integrity signaling. The analyses indicate new and diverse roles for *XTH* genes in secondary xylem cell differentiation.

## RESULTS

### *AtXTH4* and *AtXTH9* are homologs of major secondary vascular tissues XET encoding genes, *PtXTH34* and *PtXTH35*

To illustrate the importance of *XTH* genes in secondary growth, we analyzed the expression patterns of the *XTH* family members across the wood developmental zones available in the AspWood database (http://aspwood.popgenie.org; Sundell et al., 2017). Out of the recently updated census of 43 *Populus XTH* genes (Kumar et al., 2019), 26 were expressed in AspWood and the majority belonged to cluster ‘e’ (**Figure S1**), which groups genes with peak expression in the cambium and radial expansion zone, coinciding with the peak of XET activity (Bourquin et al., 2002). The subclade of *PtXTH34* (also known as *PtXET16A*) and the subclade of *PtXTH35* include the most highly expressed genes of this cluster, with documented (Kallas et al., 2005) and predicted (Bauman et al., 2007) XET activity, respectively. Arabidopsis *AtXTH4* and *AtXTH9* genes known to be highly expressed in stems and seedlings (Yokoyama and Nishitani, 2001), similar to *PtXTH34* and *PtXTH35*, respectively, were selected (**Figure 1 A**) for functional studies during secondary growth. Promoters of *AtXTH4* and *AtXTH9* were active in developing secondary vascular tissues in secondarily thickened hypocotyls and basal stems, where secondary growth occurs. The signals were observed in the vascular cambium, and adjacent developing secondary xylem and phloem, but not in the interfascicular fibers (**Figures 1 C-J**). This pattern matches the expression of their homologous clades in aspen (**Figure 1 B**), supporting their conserved functions in secondary growth in the two species.

**Figure 1.**
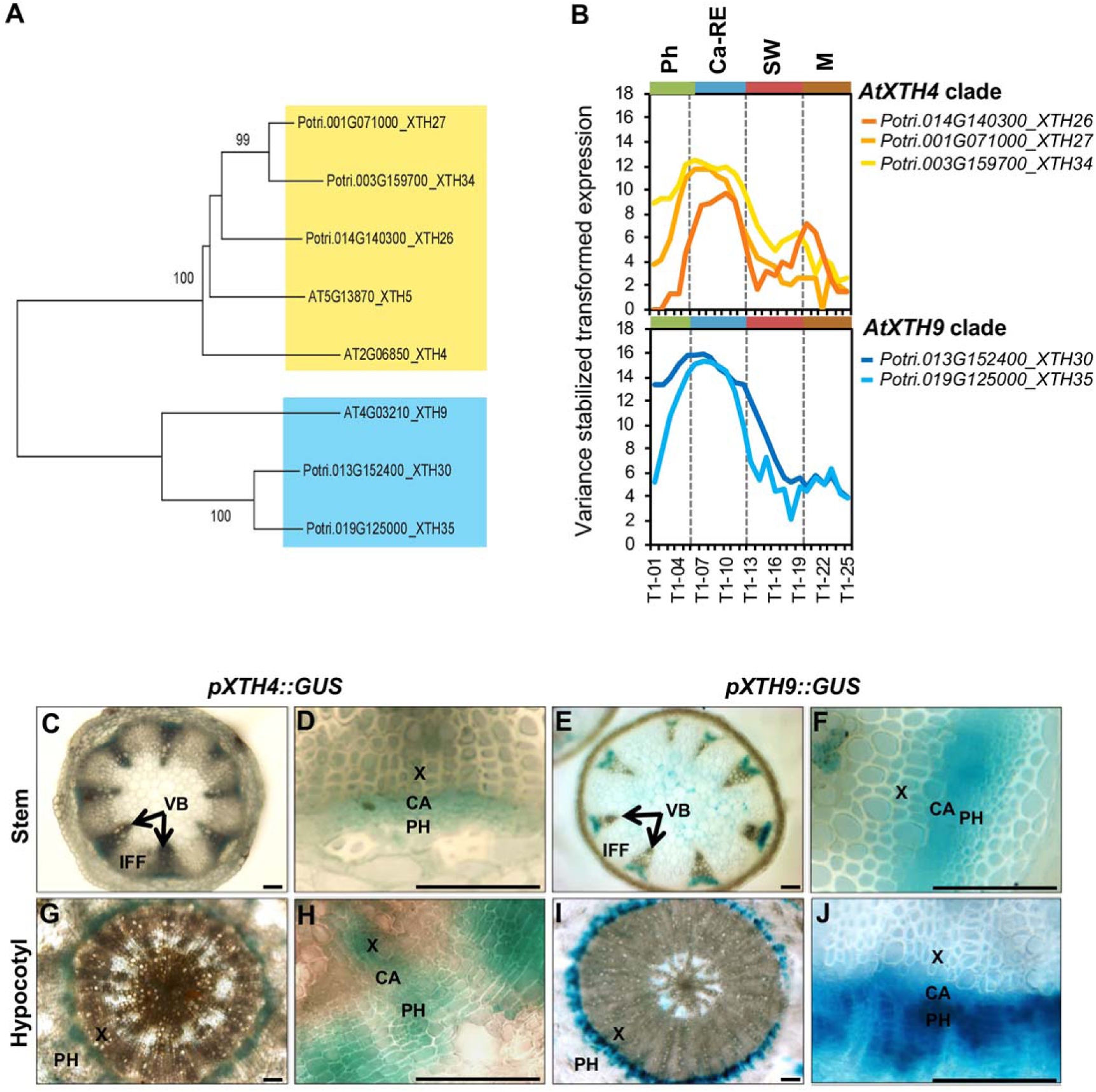
Clades *AtXTH4 and AtXTH9* in *A. thaliana* and *P. trichocarpa*. **A.** Phylogenic tree constructed using Neighbor-Joining (NJ) method of MEGA7 in default mode using MUSCLE-alligned protein sequences (http://phylogeny.lirmm.fr/phylo_cgi/index.cgi), with bootstrap test of 1000 replicates shown in % beside branches. **B.** The expression patterns of *P. trichocarpa* members of *AtXTH4* and *AtXTH9 clades* in different wood developmental zones (http://aspwood.popgenie.org). Ph-phloem, Ca-RE – cambium-radial expansion zone, SW – secondary wall formation zone, M – maturation zone. **C-J.** *AtXTH4* and *AtXTH9* promoter activity in *A. thaliana* mature inflorescence stem and hypocotyl as visualized by GUS histochemistry. In the inflorescence stems (**C-F**), the expression of both genes was detected in vascular bundles whereas interfascicular fibers did not show any expression **(C, E)**; the close up vascular bundles **(D, F)** show signals in the vascular cambium, developing xylem, and developing and differentiated phloem. In hypocotyl (**G-J**), both genes were expressed in the region of secondary vascular tissue formation **(G, I)**, encompassing the vascular cambium, developing secondary xylem and phloem, and recently differentiated phloem cells **(H, J)**. CA, vascular cambium; X, xylem; PH, phloem; VB, vascular bundle; IFF, interfascicular fibers. Scale bars 50 µm.

### Loss- and gain-of-function XET lines show altered growth and XG signals

Lines with T-DNA insertions in *AtXTH4* and *AtXTH9* genes (**Figure 2 A**) obtained from Nottingham Arabidopsis Stock Centre (NASC) were purified by repeated back-crossing until single inserts with segregation ratio 3:1 were obtained. RT-PCR analysis in basal inflorescence stems and secondarily thickened hypocotyls of flowering plants detected no transcripts of *AtXTH4* in *xht4* or those of *AtXTH9* in *xth9*. The mutants were crossed to generate the double mutant *xth4x9*. As expected, there was no expression of *AtXTH4* in the double mutant, however a low level of residual expression of *AtXTH9* was detected in this genetic background in the hypocotyl, indicating that the *xth9* mutant was conditionally slightly leaky. There was also a compensatory upregulation of *AtXTH9* and *AtXTH4* in single *xth4* and *xth9* mutants, respectively. For gain of function studies, the highly expressing line 18 (OE18) with the coding sequence of hybrid aspen (*P. tremula* L. x *tremuloides* Michx.) *PtxtXTH34* (AF515607) under the control of 35S promoter (Miedes et al., 2013) was used.

**Figure 2.**
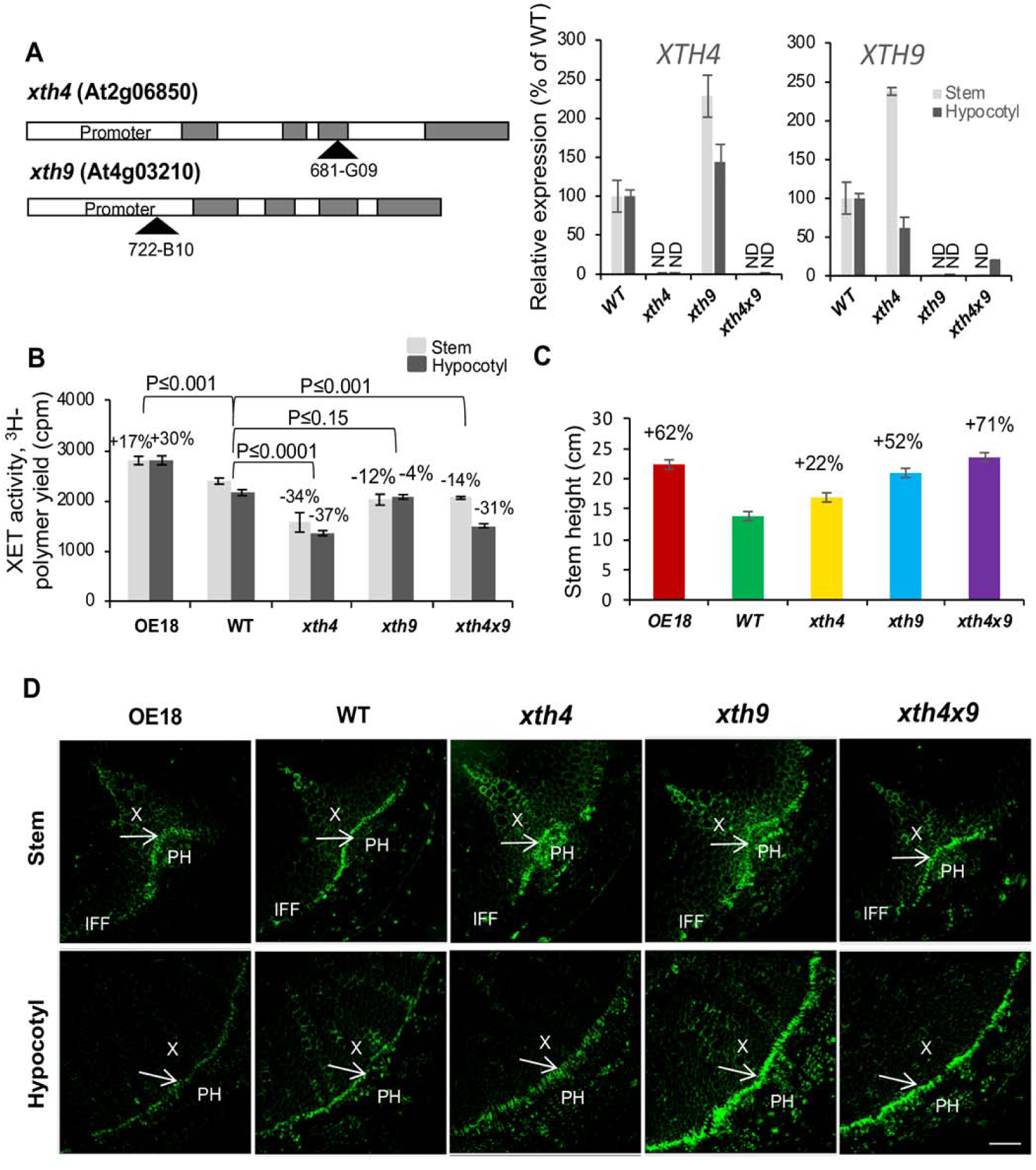
Mutations in *AtXTH4* and *AtXTH9* affect XET activity in stem and hypocotyl, stem height, and XG signals in secondary vascular tissues. A. Positions of T-DNA inserts in *AtXTH4* and *AtXTH9* genes, and expression of target genes in mutant single insert lines and in the double mutant analyzed in inflorescence stem base and in secondarily thickened hypocotyls by qRT-PCR. ND – not detected.. B. XET activity in protein extracts from basal part of inflorescence stems and in hypocotyls of 6 weeks old *xth4* and *xth9* mutants, double mutant *xth4x9*, and overexpressing line OE18 determined by the incorporation of ^3^H-labeled XG oligosaccharide (XGO) acceptor to insoluble residue and normalized to tissue fresh weight. **C.** Stem height in 6-weeks old plants grown in long day conditions. In B, *P* values for the post-ANOVA t-test are shown, and in **C** - significant differences from the WT are shown by % change (post ANOVA t-test, P≤5%), *N*=5 and 30, respectively, means ± SE. **D.** Immunolocalization of XG with CCRC-M1 antibody in vascular tissues of basal stem and hypocotyl of 6 weeks old plants. Arrows point to cambium; X, xylem; PH, phloem; IFF; interfascicular fibers. Scale bar = 50 µm.

XET activity in protein extracts from stems and hypocotyls was reduced in *xth4* and *xth4x9,* not significantly affected in *xth9* mutant and increased in OE18 line as compared to the wild type (WT) (**Figure 2 B**). Small impact of the single mutations and overexpression on activity in the protein extract was not surprising considering that many other *XTH* family genes could be contributing to the measured activity, and that *xth9* mutantion was conditionally leaky, and *AtXTH4* and *AtXTH9* exhibited compensatory reciprocal transcript activation in single mutant background (**Figure 2A**). Despite the modest changes in XET activity of extracts, the lines exhibited clearly detectable phenotypical changes as compared to WT. Stem height was stimulated in both single and in double mutants, as well as in overexpressing plants (**Figure 2 C**). Second, there were clear changes in cell wall XG signals from the monoclonal antibody CCRC-M1 in the cambium region tissues, as detected by immunofluorescence. The signals were decreased in the OE18 line and increased in *xth9* mutant in stem and hypocotyl sections, and to a small degree increased in the double mutant *xth4x9* in the hypocotyl and in *xth4* mutant in the stem (**Figure 2D**), suggesting that XET activity either affects the amount of primary walled tissues or the content of XG epitopes in these tissues, with defects in *AtXTH4* and *AtXTH9* being non-additive.

### *AtXTH4* and At*XTH9* mediate xylem cell expansion and inhibit secondary xylem production

We have previously reported that overexpression of *PtxtXTH34* in hybrid aspen increased vessel diameter (Nishikubo et al. 2011). To study the role of *AtXTH4* and *AtXTH9* in xylem cell expansion, the sizes of secondary xylem cells were measured in hypocotyls of 6 weeks old *xth4*, *xth9* and *xth4x9* mutants and OE18 plants. Vessel element diameter was reduced in the mutants, and increased in OE18 (**Figure 3A**). Vessels element length was also reduced in all three mutants, and the reduction was primarily caused by the shorted tails. This is interesting, since the length of tails partially depend on the placement of the perforation plate and partially on the tail intrusive growth. Therefore, we have analyzed the length of fibers that uniquely elongate by intrusive growth and found reductions in all three mutants but no marked effect was observed in OE line. This suggests that both genes, *AtXTH4* and *AtXTH9*, are positively regulating intrusive xylem cell elongation and vessel expansion. To our knowledge, this is the first report implicating XET in intrusive growth. Fiber diameters, which enlarge by symplastic expansion, were not consistently affected by XET activity, being enlarged in both the OE18 line and in the double mutant (**Figure 3 A**).

**Figure 3.**
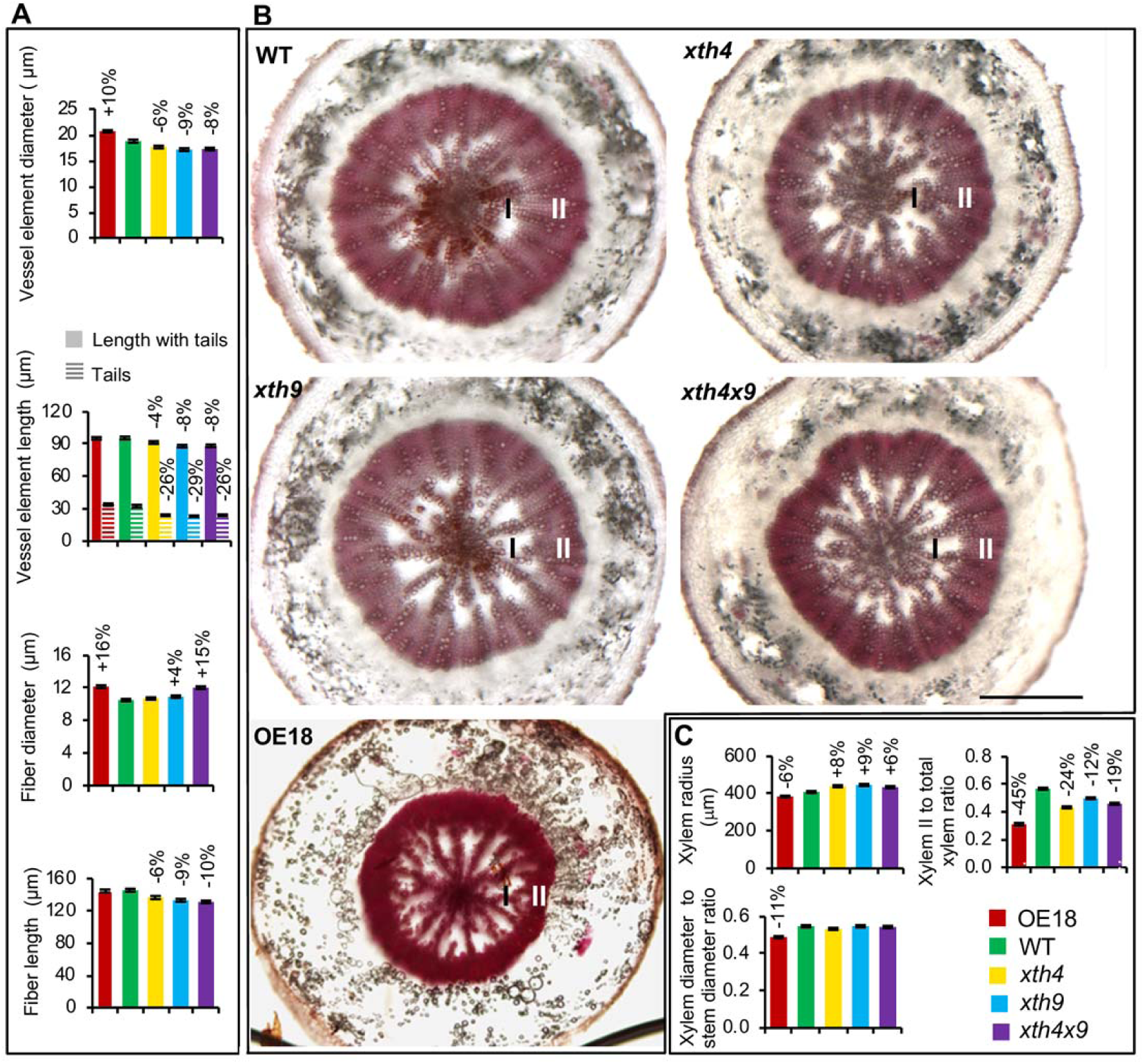
Effects of altered XET expression on secondary xylem anatomy in hypocotyls of 6 weeks old *A. thaliana* plants. **A**. Sizes of secondary xylem cells measured in wood macerates from 10 plants. **B.** Light microscopy images of hypocotyl cross sections stained with phloroglucinol. I - xylem I; II - xylem II. Scale bar = 500 µm. **C.** Morphometric analysis based on sections of 10 plants. Secondary xylem radius, xylem diameter to stem diameter ratio, and the ratio of xylem II radial width to total xylem radius. In **A** and **C** - statistically significant differences from the WT are shown by % change compared to WT (post ANOVA Student’s t-test, P≤5%), means ± *SE*.

Anatomy of secondarily thickened hypocotyls revealed changes in xylem radius; it was stimulated in the mutants and inhibited in the OE18 line (**Figure 3 B, C**). The radius reflects the number of cells in radial files and their diameters. Since in Arabidopsis hypocotyl, vessels tend to be arranged in radial files, some of which extend from the cambium to the pith (**Figure 3 B**), and the diameters of vessel elements were decreased in the mutants and increased in the OE18 line, it can be concluded that the number of xylem cells per radial file must have been increased in the mutants and decreased in the OE plants. This implicates XET in negative control of periclinal cell division in xylem mother cells. This conclusion is further supported by the reduced ratio of secondary xylem diameter to total stem diameter in OE plants (**Figure 3 C**).

Moreover, the secondary xylem development was affected in the analyzed genotypes. Secondary xylem development is known to have two distinct growth phases; the early phase, when only vessel elements and primary walled parenchyma cells are produced (xylem I), and the later stage, when the differentiation of secondary walled fibers occurs (xylem II). The ratio of xylem II to total xylem was reduced in both mutants and OE plants compared to WT, with the strongest reduction (by 45%) observed in the OE18 line (**Figure 3 B, C**). This result implicates XET in the control of transition between xylem I and xylem II phases, although these responses did not follow XET activity consistently.

### XET affects cell wall thickness and ultrastructure in mature xylem cells

To test if XET has any influence on xylem cell wall ultrastructure, we studied secondarily thickened hypocotyls and stem bases by transmission electron microscopy (TEM). In the woody plants, the secondary wall typically includes three layers, S1, S2, and S3, which can be observed by TEM (Mellerowicz et al., 2001). Similar ultrastructure was reported for *A. thaliana* interfascicular fibers (Zhong and Ye, 2015). We have also observed mostly three partite secondary walls in WT *A. thaliana* plants, in the fibers and vessel elements of stems and hypocotyls as well as in the interfascicular fibers (**Figures 4 A, B; S2-S4**). Unexpectedly, the XET modified plants exhibited severe alternations in the number of secondary cell wall layers in these cell types. In all types of fibers (hypocotyl secondary xylem fibers, stem fascicular fibers, and stem interfascicular fibers), typically only two secondary wall layers were observed in *xth4* mutant, whereas many more layers than three (up to eleven) were present in *xth9* mutant (**Figures 4 A, B; S2**). In the double mutant *xth4x9,* the number of secondary wall layers was intermediate between those observed in *xth4* and *xth9*. The overexpressing line had unchanged or reduced number of layers in the hypocotyl, and increased number of layers in the stem, compared to WT (**Figure 4 B)**. Increased cell wall layer number was observed in the vessel elements of the *xth9* mutant as well, in both hypocotyls and stems, but the change was much less pronounced than in the fibers (**Figures S3, S4**).

**Figure 4.**
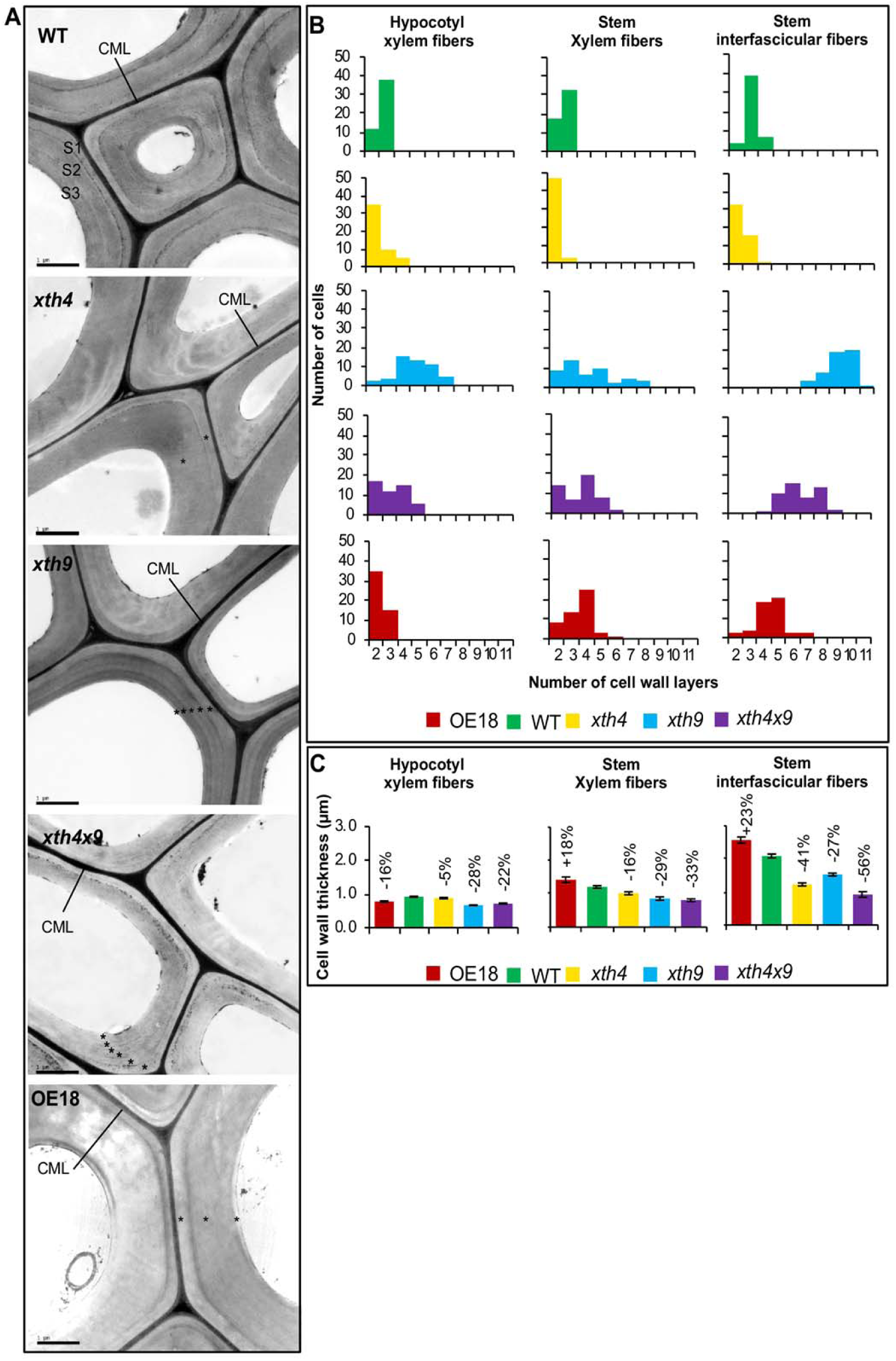
Effect of altered XET levels on secondary cell wall morphology of fibers as visualized by transmission electron microscopy (TEM) in 8 weeks old plants. **A.** TEM micrographs of hypocotyl secondary xylem fibers showing cell wall ultrastructure. Scale bar = 1 μm; CML - compound middle lamella; S1, S2, S3 – secondary cell wall layers. Note the abnormal secondary wall layering in the mutants and OE plants shown by asterisks. **B.** Frequency distributions of cells with different number of cell wall layers in hypocotyl and stem xylem fibers, and interfascicular stem fibers. 25 cells were scored in each of two plants. **C.** Secondary cell wall thickness in the different types of fibers. Statistically significant differences from the WT are shown by % change above the bars (post ANOVA Student’s t-test, P≤5%, *N* = 100 for hypocotyl, 15 for stem fibers, means ± *SE*.

Cell wall thickness in the hypocotyl and stem fibers and vessel elements was reduced in all the mutants, whereas in the overexpressing line it was either reduced (in hypocotyl fibers and stem vessel elements) or increased (in stem fibers) (**Figures 4 C; S3, S4**).

### XET affects cell wall composition in basal inflorescence stems

To test if there was any change in chemical composition of cell walls correlating with the morphological changes observed in xylem cells, alcohol-insoluble residue (AIR1) was prepared from the basal segments of inflorescence stems and analyzed by several techniques.

Pyrolysis-MS analysis detected substantial increases in lignin (total – by up to 53%, G – by up to 58%, and S – by up to 117%) in all three mutants, with most prominent effects in the double mutant *xth4x9*, whereas no difference compared to WT was found in overexpressing plants (**Figure 5 A**). Consequently there was a significant increase in S/G ratio in all mutants, the most prominent in the *xth9* (by 43%). Concomitantly, significant decreases in carbohydrate to lignin ratios were evident in all mutants by up to −40% in *xth4x9*.

**Figure 5.**
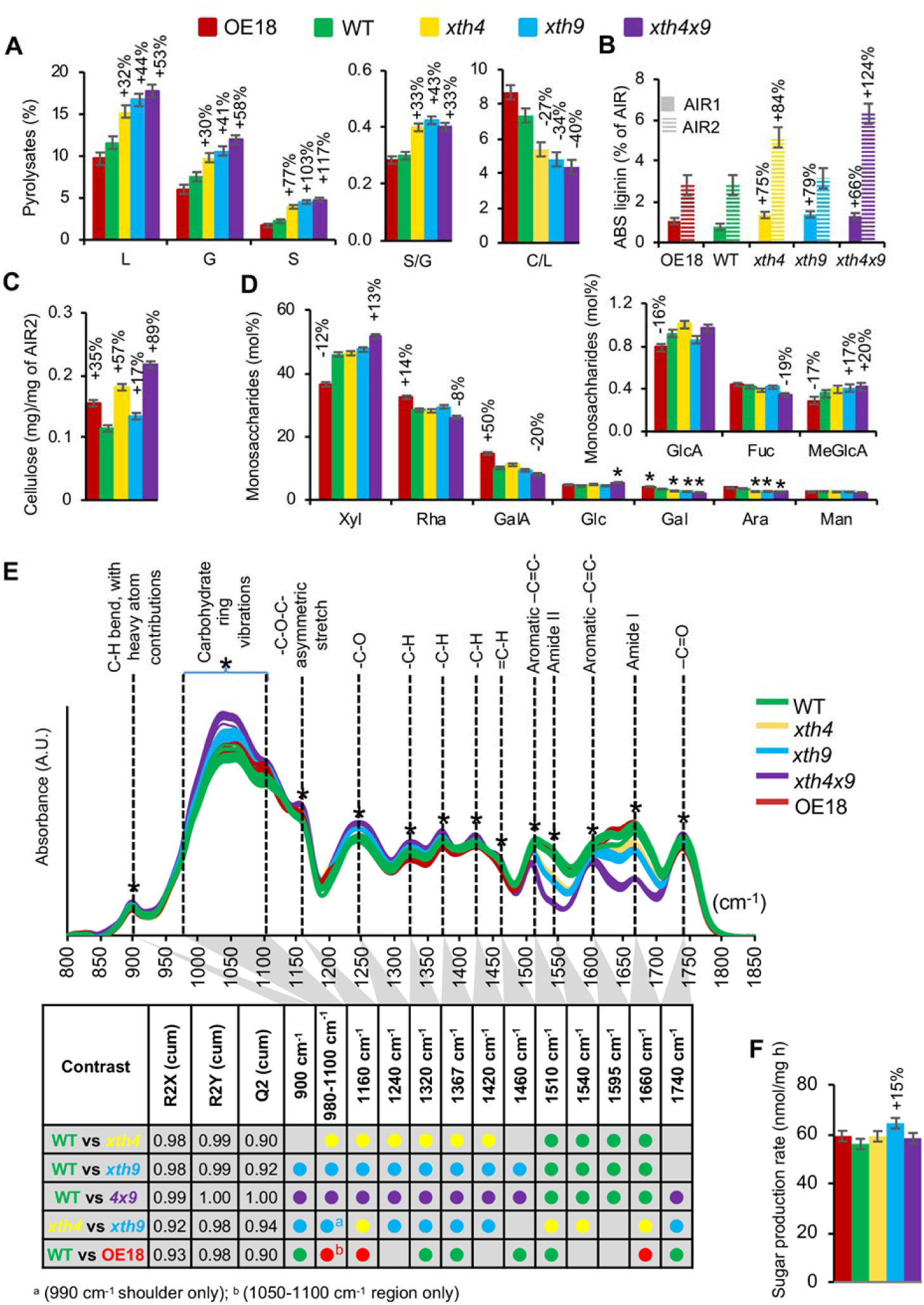
XET affects cell wall chemistry in basal inflorescence stem. **A.** Pyrolysis GC/MS of alcohol insoluble residue (AIR1). L-total lignin, G-guaiacyl lignin, S-syringyl lignin, C-carbohydrates. **B.** Acetyl bromide-soluble (ABS) lignin in AIR1 before (AIR1) and after (AIR2) amylase treatment. **C.** Updegraff cellulose content in AIR2. **D.** Monosaccharide composition of AIR2 determined by methanolysis and trimethylsilyl (TMS) derivatization. **E.** Diffuse reflectance Fourier-transform infrared (FTIR) spectra of AIR1. Asterisks denote bands that are significantly different (≥50% correlation) in the different contrasts shown in the table, according to OPLS-DA (orthogonal projections to latent structures – discriminant analysis) models using 1+2 (predictive + orthogonal) components. Coloured dots indicate genotypes with significantly higher signals. Score plots and loadings for the models are shown in Figure S5. **F.** Sugar yields of dried stems in saccharification analysis after alkaline pretreatment. Data are means ± *SE*, *N*=3 for A-D or 5 for F, percentage changes or asterisks are given for means significantly different from WT (post-ANOVA t-test, P ≤ 0.05).

Acetyl bromide soluble (ABS) lignin content was determined in AIR1 directly, as well as after the amylase treatment (AIR2). ABS lignin was substantially increased in AIR1 in all the mutants (by up to 79% in *xth9*) while it was unaffected in the OE line, in agreement with pyrolysis-MS results (**Figure 5 B**). However, lignin amounts in the de-starched samples increased only in *xth4* and *xth4x9* mutants, but not in *xth9*, which was most affected in the AIR1 fraction. This indicates that the *xth9* mutant either contained less starch in AIR1 than other lines (contributing to artefactually high ABS lignin content when expressed per AIR1 weight), or that the lignin induced by *xth9* mutation is vulnerable to amylase treatment (for example by containing relatively more water soluble components).

Crystalline cellulose content in the amylase treated alcohol insoluble residue (AIR2) was increased in the double mutant (by 89%), and to a lesser degree in the single mutants (by 57% in *xth4* and 17% in *xth9*) (**Figure 5 C**). Interestingly, the OE18 line also had higher cellulose content (by 35%) than the wild type. Matrix polysaccharide composition analysis of AIR2 revealed increased Xyl and 4-*O*-methyl glucuronic acid (Me-GlcA) in the double mutant *xth4x9*, indicative of increased glucuronoxylan content, and decreased rhamnose (Rha), GalA, Gal and arabinose (Ara), indicative of reduced pectin content (**Figure 5 D**). Opposite changes were observed in OE18 line, indicative of decreased glucuronoxylan and increased pectin content as compared to wild type. Taken together, these changes suggested that there could be a change in relative contributions from primary and secondary walls in the basal stems of double mutant and the OE18 line, the former having more secondary walls, and the latter more primary walls than the wild type.

To reveal other changes in cell walls, diffuse reflectance Fourier-transform infrared (FTIR) spectra of AIR1 were compared among the genotypes revealing clear differences (**Figures 5 E, S5**). The most affected peak was 1660 cm^-1^, where signals were substantially reduced in the *xth* mutants, with the largest decrease in *xth4x9*, a larger decrease in *xth9* than in *xth4*, and significant increase in OE18. The 1660 cm^-1^ band corresponds to the amide I vibration of proteins and as such can be connected to XET proteins covalently attached to XG in cell wall. The proteinic origin of this band was confirmed by similar trends observed for the 1540 cm^-1^ band (amide II). Bands associated to aromatic -C=C-skeletal vibrations (ca. 1510 and 1595 cm^-1^), which are typically interpreted as lignin signals, were also affected, but these bands overlap with the amide bands. Thus, they are highly susceptible to strong changes in the amide bands observed in the mutants and OE plants. This interpretation is consistent with the wet chemistry data obtained from the same material (**Figure 5 A, B**). The mutants also showed a significant increase in bands traditionally associated to polysaccharidic compounds (e.g. carbohydrate ring vibrations between 980-1100 cm^-1^, -C-O-C-asymmetrical stretch at 1160 cm^-1^, and diverse -C-H related vibrations at 1320 and 1420 cm^-1^), indicating a higher proportion of carbohydrates, potentially cellulose (1160 cm^-1^; Dokken et al., 2005) and xylan (1320 and 1420 cm^-1^; Kačuráková et al., 1999), as compared to the wild type. The cellulose associateed -C-O-C-band (1160 cm^-1^) also increased in the OE18 line compared to WT, and in *xth4* compared to *xth9* (**Figures 5 E, S5**). These carbohydrate-related signals were consistent with the wet chemistry data (**Figures 5 C, D**).

Considering substantial alterations in cell wall chemistry in the basal stems and in the ultrastructure of secondary walls in the studied *xth* mutants and OE plants, we examined their saccharification characteristics. However, sugar production rate from the dried stems after alkaline pretreatment (Gomez et al., 2010) only improved in the *xth9* mutant (**Figure 5 F**).

### *In situ* FTIR microspectroscopy data reveal chemical differences in secondary walls of *xth* mutants and OE plants

The changes in cell wall composition in the *xth* mutants and OE plants revealed by the bulk analyses of basal inflorescence stem tissues suggested relative changes in the amount of primary and secondary walls. To reveal any potential chemical changes specifically in secondary walls, we carried out *in situ* FTIR microspectroscopy analysis of fascicular and interfascicular fibers, which are the two cell types highly contributing secondary cell wall material in the basal inflorescence stem. In addition, we have analyzed xylem fibers in hypocotyls to verify if similar changes are induced in this organ.

*In situ* FTIR spectra of either stem or hypocotyl fiber walls did not reveal difference in *xth4* compared to WT (**Figure S6 A-D**). In contrast, *xth9* mutant showed differences in signal intensities of several bands, notably there was an increase in the 1510 cm^-1^, and a decrease in the 1320 cm^-1^ signals in both analysed organs. The increased intensity of the 1510 cm^-1^ band, which was also seen in hypocotyl in the double mutant, and which was observed in *xth9* compared to *xth4*, could be related to a more cross-linked form of lignin (Faix, 1991). Several bands were altered in the OE18 plants, most of them being consistent between stems and hypocotyls (**Figure S6 A-D**). Beside the 1510 cm^-1^ (indicative of aromatics, although influence of the neighbouring amide II band cannot excluded) and 1460 cm^-1^ bands (observed for both lignin and hemicellulose, potentially originating from =C-H functionalities) that had increased proportional intensities, we observed an increased intensity of the 1240 cm^-1^ (-C-O) and 1420 cm^-1^ bands (-C-H), and a shift of the 1740 cm^-1^ band (-C=O) to higher wavenumbers (indicating higher energy vibrations of this functional group). Together, the increase of -C-H, -C-O band intensities and the shifted -C=O band may indicate an increased level of esterification. Conclusively, *in situ* microspectroscopy FTIR revealed clear chemical changes in the chemical composition of mature fiber walls.

### *In situ* Raman microspectroscopy data reveal differences in lignin in secondary walled cells of *xth* mutants and OE plants

To obtain more support for the chemical changes in cell walls of mature fibers, we carried out *in situ* Raman microspectroscopy, which provides a higher spatial resolution and spectra with narrower bands and significantly reduced contribution from hemicelluloses and pectins compared to FTIR spectroscopy. The spectra of interfascicular fibers, where neither *AtXTH4* nor *AtXTH9* were expressed, and fascicular fibers, where both these genes were found expressed (**Figure 1 C-J**), were taken separately to address the question of cell-autonomous *versus* non-cell autonomous effects. Pairwise comparisons by cell type, between each genotype and WT, revealed significant differences among genotypes (**Figure S7, A-D**). Considering only the differences between genotypes that were consistent between at least two cell types, we noted higher signals in *xth4* compared to WT from several bands corresponding to lignin, such as 595 cm^-1^, 845 cm^-1^, and 1270 cm^-1^ or xylan and lignin – 1250 cm^-1^ (**Figure 6 A, B**). In contrast, bands corresponding to cellulose were reduced in fiber cell walls in this genotype (900, 970 and 1000 cm^-1^) together with bands corresponding to non-cellulosic polysaccharides, cellulose or lignin (1330, 1365, and 1390 cm^-1^). Several of these bands also appeared similarly affected in other mutants, but only in one tissue type. Higher signals were observed in the *xth9* mutant compared to WT for signals around 1215 cm^-1^, corresponding to xylan and lignin, and at 1505 cm^-1^, corresponding to aromatics (lignin). These bands showed increased intensities in *xth9* than *xth4* but only in one of the studied tissues. Other lignin-related bands at 725 and 1655 cm ^-1^ were increased in OE18. This result supports the conclusions of an altered lignin structure / composition in the fibers in the *xth* mutants and OE plants, compared to WT, and indicates that distinct lignin alterations have occurred in *xth4*, *xth9*, and in OE18. Intruigungly, the changes were most frequently observed in interfascicular fibers, suggesting their non-cell autonomous induction.

**Figure 6.**
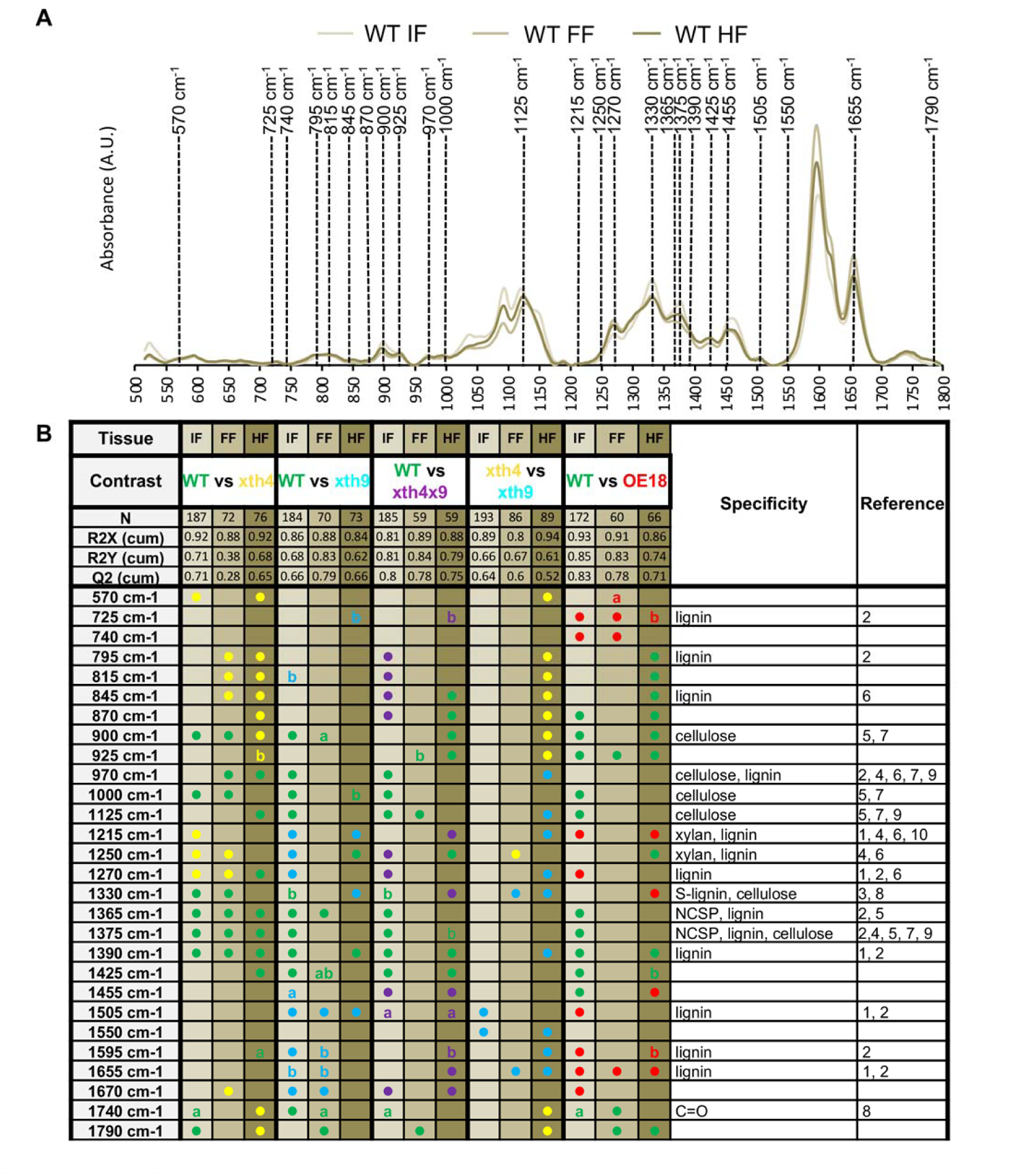
*In situ* Raman microspectroscopy of cell walls in fibers in three different tissues in *xth* mutants and overexpressing plants. **A.** Representation of average WT spectra of interfascicular fibers (IF); fascicular fibers (FF) and hypocotyl fibers in secondary xylem (HF), with the bands that were significantly contributing to the separations in at least two of the studied tissues in any of the constructs shown in **B**. **B.** Summary of OPLS-DA models of pairwise comparisons using 1+2 (predictive + orthogonal) components. Coloured dots indicate genotypes with higher levels of given signals. Details of the models are given in **Fig S7.** References: (1) Agarwal, 1999; (2) Agarwal et al., 2011; (3) Agarwal and Atalla, 2010; (4) Agarwal and Ralf, 1997; (5) Kačuráková et al., 1999; (6) Larsen and Barsberg, 2010; (7) Schenzel and Fischer, 2001; (8) Schulz and Baranska, 2007, (9) - Wiley and Atalla, 1987; (10) Zeng, 2016; NCSP – non-cellulosic, structural polysaccharides; a - low end, i.e. redshift; b - high end, i.e. blueshift.

To asses if any chemical change could be related to the layering phenotype of secondary walls, we used the cell wall layers as Y (response) variables in an OPLS model of Raman spectra. The average number of cell wall layers were obtained from TEM analyses for interfascicular, fascicular, and hypocotyl fibers in each genotype. The resulting scatter plot (**Figure 7 A**) shows that the predictive component could identify spectral changes correlated to the number of cell wall layers, while the first orthogonal component was mainly related to tissue type. Thus, there were specific spectral features that were correlated to the numbers of secondary wall layers, irrespective of tissue type. The corresponding Loadings (**Figure 7 B**) identified these features as bands at 900, 1040, 1150, 1330, 1460, and 1505 cm^-1^, correlated with the high layer number, and bands at 1270 and 1565-1570 cm^-1^, correlated to low number of layers. The band at 900 cm^-1^ (correlated to high numbers of cell wall layers) can be found in the spectra of all major polysaccharidic polymers, with cellulose and hemicellulose among the most common associations (Gierlinger, 2018). The band at 1040 cm^-1^ is located firmly in the region that is dominated by carbohydrate ring vibrations often used to assess their proportions (Gierlinger, 2018). The band at 1150 cm^-1^ is usually attributed to -C-O-C-vibrations but it is rather unspecific, being present in both polysaccharidic but also lignin polymers. The bands at 1330 and 1505 cm^-1^, on the other hand, while also correlated to high number of layers, have been assigned to different lignins (Gierlinger, 2018). The band at 1460 cm^-1^ (correlated to high numbers of cell wall layers) is attributed to -C-H, and the one at 1270 cm^-1^ (correlated to low numbers of cell wall layers) to -C-O vibrations, but they are seldom specific. The shoulder between 1565 – 1570 cm^-1^ (correlated to low number of layers) is in the spectral region of C=X vibrations, where X is either another carbon (often aromatic, making it a lignin-related band) or an oxygen. Both excludes the possibility of this band originating from cellulose. Thus, bands that can be assigned to cellulose are found correlated to a higher number of cell wall layers, while lignin-associated bands are found correlated to both higher and lower numbers of cell wall layers, indicating an altered lignin composition. Overall, these data indicate specific changes in lignin and cellulose in secondary walls as a function of the number of secondary wall layers, irrespective of cell type.

**Figure 7.**
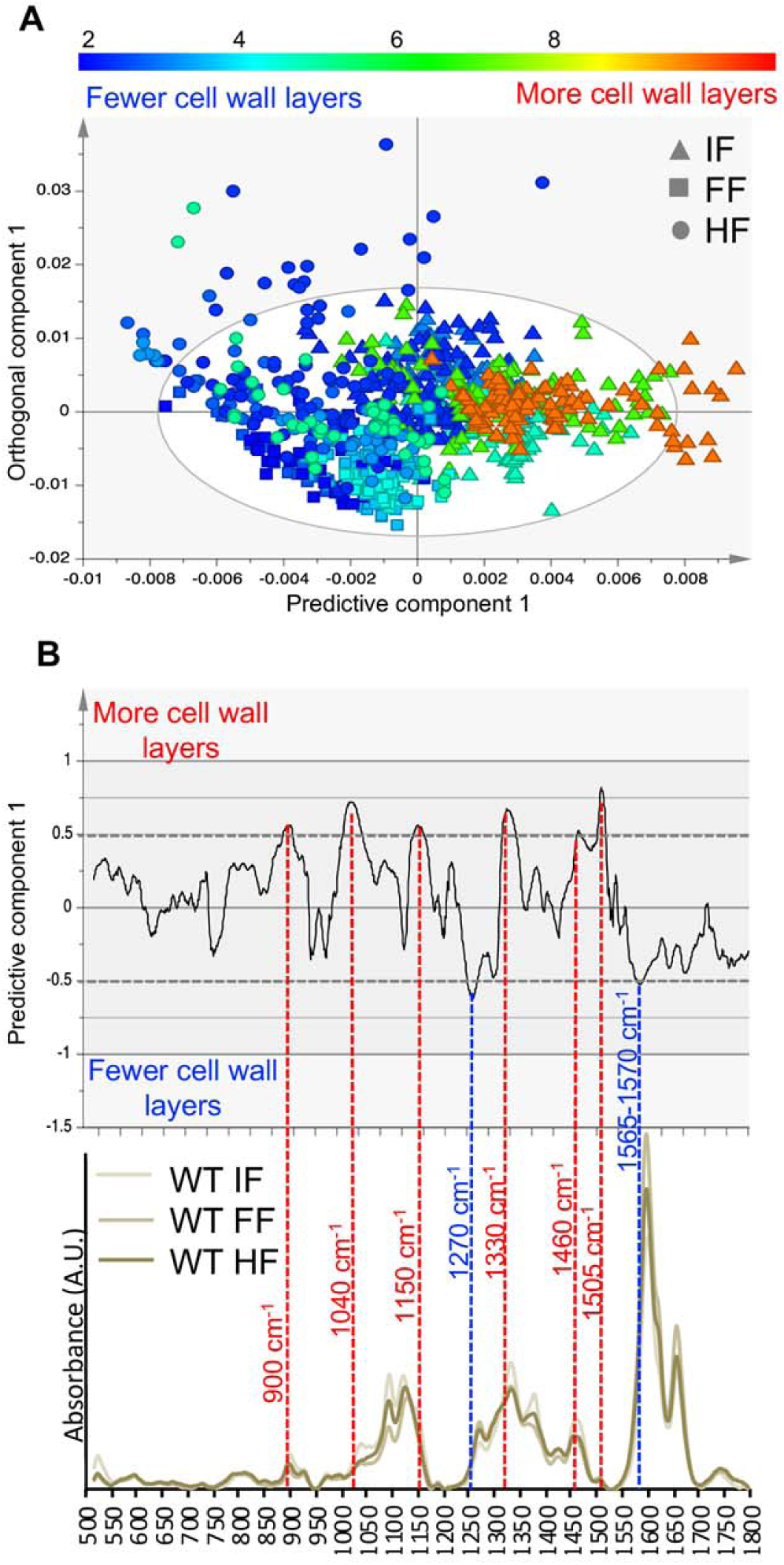
OPLS model correlating average number of cell wall layers to Raman microspectroscopic features in three tissues: interfascicular fibers (IF), fascicular fibers (FF), and hypocotyl secondary xylem fibers (HF) **A.** Scatter plot showing separation according to the number of cell wall layers (predictive component) and tissue type (orthogonal component). **B.** The corresponding Loadings Plot and representative WT spectra for the three tissues. The spectral bands with more than 50% correlation levels (red dashed lines) are marked by band numbers. Bands at 900 cm^-1^, 1040 cm^-1^, 1150 cm^-1^, 1330 cm^-1^, 1460 cm^-1^, 1505 cm^-1^ are proportionally more intensive when there are more cell wall layers, whereas bands at 1270 cm^-1^ and a shoulder 1565-1570 cm^-1^ are proportionally more intensive when there are fewer cell wall layers. The model details are as follows: 820 spectra, 1+3 (predictive + orthogonal) components, R2X(cum) = 0.928, R2Y(cum) = 0.383, Q2(cum) = 0.377.

### Effect of altered *AtXTH4* and *AtXTH9* on gene expression in lignin and cellulose biosynthesis, and cell wall integrity sensing pathways

To test if the reported changes in secondary walls in *xth* mutants and OE plants were induced indirectly by activation of biosynthetic and signaling pathways, we analyzed transcripts of selected sets of genes related to secondary wall biosynthesis and to cell wall integrity sensing pathways in the basal part of inflorescence stems.

Among the genes representing six enzymatic activities essential for monolignol biosynthesis, we detected strong upregulation in the *xth* mutants in genes of the early pathway, *PHENYLALANINE AMMONIA-LYASE 1* (*PAL1*) and *PAL4*, downregulation in all mutants in *4-COUMARATE:COENZYME A LIGASE* (*4CL*), and the upregulation of genes representing the late pathway, *p-COUMARATE 3-HYDROXYLASE* (*C3H*), *CAFFEOYL-CoA O-METHYLTRANSFERASE 1* (*CCoAOMT1*) and *FERULATE 5-HYDROXYLASE* (*F5H)* (**Figure 8 A**). *PAL1*, *PAL4*, *C3H* and *F5H* were especially highly upregulated in *xth9* mutant. The upregulation of these monolignol biosynthetic genes could explain increase in lignin content in the mutants and a distinct lignin composition in *xth9* detected by pyrolysis and wet chemistry analyses (**Figure 5 A**). The OE plants, on the other hand, exhibited many lignin biosynthesis genes downregulated, including *PAL2*, *PAL4*, *CINNAMATE 4-HYDROLASE* (*C4H*), *4CL*, *C3H* and *F5H*.

**Figure 8.**
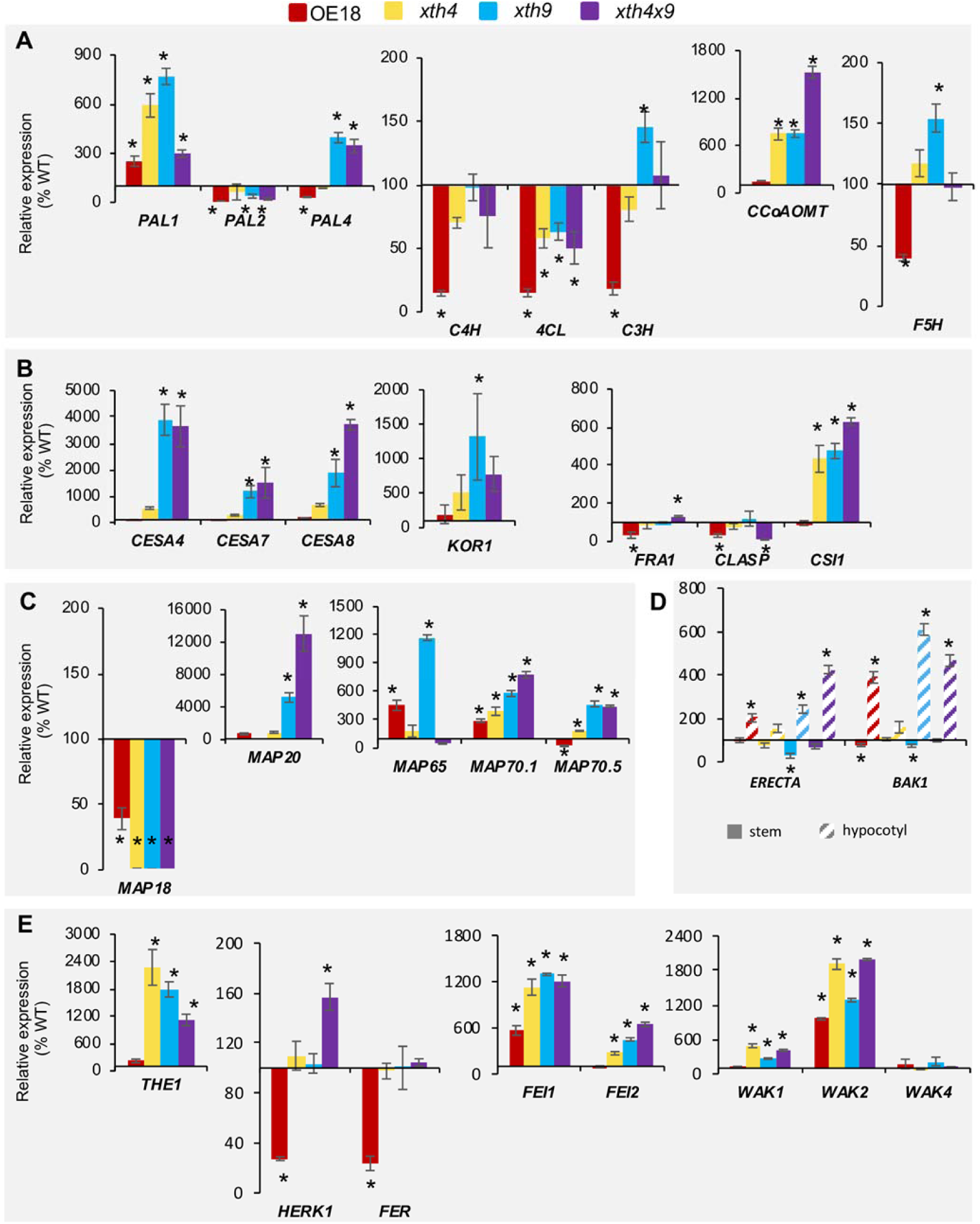
Expression levels of genes affected in the *xth* mutants and overexpressing plants. Genes related to lignin biosynthesis **(A)**, secondary wall cellulose biosynthesis **(B)**, microtubule dynamics **(C)**, secondary xylem regulation in hypocotyl **(D)**, and cell wall integrity sensing **(E)** were determined by quantitative RT-PCR. Expression levels were normalized to WT (100%). Data are means ± *SE*, *N*=3 biological replicates, asterisks indicate means significantly different from WT (post-ANOVA t-test, *P* ≤ 0.05).

The genes encoding cellulose synthase complex proteins such as secondary wall CESAs (*CESA4*, *CESA7* and *CESA8*), *KORRIGAN1* (*KOR1*), and *CELLULOSE SYNTHASE INTERACTING1* (*CSI1*) were greatly upregulated in *xth9* and *xth4x9* mutants, and in the latter case, also in *xth4* mutant (**Figure 8 B**). The *FRAGILE FIBER 1 (FRA1*) encoding a kinesin like protein, regulating cellulose microfibril angle (Zhong et al., 2002) and CLIP170-ASSOCIATED PROTEIN (CLASP) that promotes microtubule stability (Ambrose et al., 2007) showed some variability in *xth4x9* mutant and mild downregulation in OE line. Similarly, four genes encoding microtubule-binding proteins with known function in secondary wall cellulose biosynthesis (Korolev et al., 2007; Rajangam et al., 2008; Pesquet et al., 2010) were highly upregulated in the mutants, especially *MAP65* in *xth9, MAP20* in *xth9* and *xth4x9,* and *MAP70.1* and *MAP70.5* in all mutants (**Figure 8 C**). Some variability was also observed in OE18 line in *MAP65*, *MAP70*.1, and *MAP70.5.* In contrast, *MAP18* encoding a MT destabilizing protein regulating cell growth in expanding cells (Wang et al., 2007) was downregulated in all mutants and OE plants.

One way these changes in biosynthetic secondary wall related genes could be induced is by the triggering of the cell wall integrity pathway. We found several key players in cell wall integrity sensing affected in the mutants and OE plants. Notably, the genes encoding plasma membrane localized receptor-like kinases (RLK) such as *THESEUS1* (*THE1*), *HERKULES 1* (*HERK1*), *FERONIA* (*FER*), *WALL-ASSOCIATED KINASES* (*WAKs*) (Li et al., 2016), and ERECTA-related LRKs *FEI1* and *FEI2* (Xu et al., 2008) were affected. *THE1* was significantly upregulated in all mutants, *HERK1* in *xth4x9,* and *HERK1* and *FER* were downregulated in OE18 line (**Figure 8 E**). Among tested WAKs, *WAK1* and *WAK2* were upregulated in all mutants and *WAK2* was also upregulated in OE line. *FEI1* and *FEI2* were upregulated in all mutants, and *FEI1* was upregulated in OE plants. These data indicate the stimulation of cell wall integrity pathways in the basal part of inflorescence stems by altered XET expression.

Moreover, since the cell wall integrity sensors eventually would need to affect global master regulatory switchers to affect vascular development seen in the mutants and OE plants (**Figure 3**), we tested one of such switchers, *ERECTA*, which is known as negative regulator of xylem II differentiation in hypocotyls (Ikematsu et al., 2017), and its putative co-receptor *BAK1* (Jorda et al., 2016). We found both *ERECTA* and *BAK1* upregulated in hypocotyls of OE plants and in *xth9* and *xth4x9* mutants (**Figure 8D**). This can provide a direct explanation of the suppressed xylem II differentiation in these plants.

## DISCUSSION

XETs have been implicated in plant growth and development as well as stress reactions to mechanical stimuli, wounding and cold (reviewed by Rose et al., 2002). In spite of the growing knowledge about their enzymatic activities, their mode of action in plant tissues is still being debated. XETs were observed expressed in growing and expanded vascular tissues, and were hypothesized to mediate both cell expansion and cell wall development in these tissues since over two decades (Xu et al., 1995; Antosiewicz et al., 1997; Oh et al., 1998; Bourquin et al., 2002; Dimmer et al., 2004; Jiménez et al., 2006). However, data from overexpression or from knock-out/suppression studies demonstrating the aspects of vascular development affected by XET are scarce (Matsui et al., 2005; Nishikubo et al., 2011). Here, we have analyzed secondary vascular development in mutants of two vascular-expressed XETs, *AtXTH4* and *AtXTH9*, in their double mutant *xth4x9,* and in plants ectopically overexpressing aspen *PtxtXTH34* – closely related to *AtXTH4*. We demonstrate the influence of these mutations and overexpression on different aspects of secondary growth, including secondary wall formation. The action of *AtXTH4* and *AtXTH9* genes was found overlapping in some cases, and distinct or even opposite – in others. Moreover, we identified non-cell autonomous effects of altered XET expression. Here we discuss the most prominent effects observed.

### *AtXTH4* and *AtXTH9* mediate secondary xylem cell diffuse expansion and intrusive tip growth

A previous study on transgenic aspen overexpressing *PtxtXTH34* indicated that XET positively regulates vessel radial expansion (Nishikubo et al., 2011). Our results for *A. thaliana* confirmed and extended these findings providing evidence for the positive effect of *AtXTH4* and *AtXTH9* not only on vessel element radial expansion, but also on tail elongation, and fiber intrusive tip growth (**Figure 3 A**). Fiber diameter growth, similar to vessel element radial expansion, was stimulated by XET overexpression, but in contrast to vessel elements, it was also increased in the fibers of *xth9* and *xth4x9* mutants. Since vessel elements develop precociously relative to fibers (Mellerowicz et al., 2001), it is likely that narrower vessels in the *xth9* and *xth4x9* mutants create more room for fiber radial expansion.

Whereas the role of XET in cell wall expansion by diffuse growth has been supported by many observations (Vissenberg et al., 2000; Osato et al., 2006; Shin et al., 2006; Liu et al., 2007; Ohba et al., 2011; Miedes et al., 2013), the role of these enzymes in the tip growth and intrusive growth has not yet been clarified. Whereas in some species, including Arabidopsis, XET activity and expression of some *XTH* genes has been observed in elongating by tip growth root hairs (Vissenberg et al., 2003; 2005b), no defects in root hair elongation have not been so far reported for any *XTH* mutants, possibly due to redundancy. Our results for *xth4* and *xth9* mutants provide evidence for their involvement in tip growth of fibers and vessel element tails. It is not clear whether effects of XETs are direct mediating mechanical properties of cell walls at elongating tips, or indirect. Nevertheless, XETs may be considered as targets for fiber length improvement programs in forest tree species. An example of association between *PtoXTH34* (former *PtoXET16A*) alleles and fiber length has been already reported for *Populus tomentosa* Carrière (Wang and Zhang, 2014).

### Deficiencies in *AtXTH4* and *AtXTH9* stimulate production of secondary walled tissues but reduce secondary wall thickness non-cell-autonomously

Chemical analyses of the stem bases clearly showed increased lignin, crystalline cellulose, and xylan-related monosaccharide contents in the *xth* mutants indicative of increased overall secondary wall content in the stem basal tissues (**Figure 5 A-C**). This was supported by an increase in the expression of genes involved in secondary wall biosynthesis (**Figure 8 A-C**). In the OE plants, on the other hand, changes in matrix polysaccharides suggested decreased xylan and increased pectins in this genotype (**Figure 5 D**), pointing to the overall decreased contribution of secondary walls. Consistently, several lignin biosynthetic genes appeared downregulated (**Figure 8 A**), suggesting an overall decreased lignification in the basal stem tissues. Curiously, the effects on secondary cell wall thickness were opposite: the fiber wall thickness was reduced in the *xth* mutants and increased in the OE plants (**Figures 4 C**). Thus, the stimulation of secondary wall biosynthesis in *xth* mutants, evident from bulk chemical analyses and gene expression, must have occurred at the whole tissue level by the upregulation of the number of cells with secondary cell walls, rather than by stimulation of secondary wall thickening. This interpretation of the data concerning the stem bases is supported by direct observations in hypocotyls, where the production of secondary xylem was induced in the mutants (**Figure 3 B**), with simultaneous suppression of the secondary wall thickness (**Figures 4 C; S4 B**). Thus, the effects of XET on secondary xylem production and thickening of secondary walls are independent. This conclusion is in agreement with the current models of independent regulation of meristematic activity leading to xylem production, and secondary wall thickening activity of differentiating xylem cells (Ramachandran et al., 2017). *AtXTH4* and *AtXTH9* deficiency can inhibit secondary wall thickening non-cell autonomously, i.e. outside of their expression range, and the effects are apparently additive for the two genes (**Figure 4 C**).

### *xth4* and *xth9* mutants display opposite changes in secondary wall layers

Secondary wall layering defects seen in *xth* mutants and OE plants are the most striking phenotypes reported in this study (**Figures 4, S2-S4**). These defects were evident in all cells with secondary walls, but were most apparent in the interfascicular fibers, where neither *AtXTH4* or *AtXTH9* was expressed. This indicates that the secondary wall layering is induced non-cell autonomously by *AtXTH4* and *AtXTH9* deficiency. Curiously, deficiency in *AtXTH4* resulted in reduction of secondary cell wall layers and opposite phenotype was induced by deficiency in *AtXTH9*. The intermediate effects on cell wall layering seen in the double mutant supports opposite effects of the two mutants. XET overexpression also affected secondary wall layering with different outcomes, depending on cell and organ type. Opposite effects of *AtXTH4* and *AtXTH9* deficiency on secondary cell wall layering are puzzling since their mutants had additive phenotypes in other measured parameters, like secondary wall thickening. They could be related to subtle differences in *At*XTH4 and *At*XTH9 enzymatic properties, which are currently unknown, or to differences in expression pattern in developing xylem cells as seen for their aspen orthologs (**Figure 1 B**). Differential expression is supported by separate co-expression networks of *AtXTH4* and *AtXTH9* in Arabidopsis revealed by GeneMANIA analysis (http://www.genemania.org; Warde-Farley et al., 2010), and for *PtXTH34* and *PtXTH35* clades in AspWood database (http://aspwood.popgenie.org; Sundell et al., 2017).

Secondary wall layering is normally related to changes in cellulose microfibril angle (MFA), which are matched and affected by the changes in cortical microtubules (Mellerowicz et al., 2001). Currently, it is unknown whether XET deficiency/surplus affect cortical microtubules in the studied genotypes. Although we have seen changes in expression of several genes encoding microtubule associated proteins (MAPs) that play essential role in MT dynamics, these changes may be due to overall secondary wall induction in the stems of mutants, as they paralleled expression of secondary wall *CesAs* (**Figure 8 B, C**). There are other reports suggesting a possible coupling between XTHs or XG, cortical microtubules and cellulose biosynthesis (Takeda et al., 2002; Sasidharan et al., 2014; Xiao et al., 2016; Armezzani et al., 2018), but it is unclear how this coupling mechanistically occurs.

The secondary wall layering deffects were observed in different plants species when cinnamoyl-CoA reductase (CCR) was downregulated (Ruel et al., 2002; Leplé et al., 2007). This was accompanied by reductions in non-condensed lignin and by the presence of abnormal lignin units. Our *in situ* FTIR and Raman analyses demonstrated increased lignin content and changes in signals related to aromatic components in the studied genotypes (**Figures 6; S6**), whereas the wet chemistry analyses (**Figure 5 A, B**) and gene expression data (**Figure 8 A**) revealed distinct changes in *xth9* mutants in lignin quality and biosynthesis compared to *xth4.* This could be a basis of opposite wall layering phenotypes in these mutants. The more direct proof of distinct chemistries in secondary walls in these mutants comes from the models correlating cell wall numbers and chemistry based on *in situ* Raman signals (**Figure 7**), which indicated changes both in polysaccharidic compounds (e.g. increasing cellulose proportions with higher numbers of cell wall layers) and lignin (mixed correlations to cell wall layer numbers, indicating altered lignin composition / structure). While the exact nature of these changes are hard to elucidate from the present data, it clearly points to an overall change in the cell wall matrix.

### XET role in perception of mechanical stimuli?

XETs have been implicated in various mechanoperception responses such as thigmomorphogenesis (Xu et al., 1995; Lee et al., 2005; Martin et al., 2014), gravitropism (Cui et al., 2005), and tension wood formation (Nishikubo et al., 2007; Baba et al., 2009; Kaku et al., 2009; Gerttula et al., 2015). XET activity conceivably could be involved in these responses by generating signaling XG-oligosaccharides (XGO) although their mode of action is poorly understood (York et al., 1984; McDougall and Fry, 1988; 1989; Kaida et al., 2010; González-Pérez et al., 2012, 2014). XET could also be involved in these processes by preventing a build-up of tension among polymers in newly deposited cell wall layers as has been proposed for xylan endotransglycosylase *PtxtXYN10A* (Derba-Maceluch et al., 2015). In such case, XET deficiency would lead to increase of tension in cell wall that could be perceived by plant cell as mechanical stress, thus probably acting via cell wall integrity sensing mechanisms (Li et al., 2016; Wolf 2017). Interestingly, the observed symptoms of *AtXTH4* and *AtXTH9* deficiency – stimulation of xylem mother cell division, suppression of secondary wall thickening in xylem cells, and increased height growth mimic those observed in aspen when secondary wall xylan synthase members *Pt*GT43B and *Pt*GT43C were suppressed (Ratke et al., 2018). Moreover, the altered expression of several receptor-like kinases, including *THE1* and *WAK2*, known to mediate cell wall integrity sensing observed in *xth9*, *xth4,* and *xth4x9* mutants as well as in XET OE plants strongly supports the modulation of the signaling via a cell wall integrity sensing pathway in these genotypes (**Figure 8E**). Upregulation of some of these receptors, for example WAK2, in both, mutants and OE plants, could provide explanation how XET deficiency and excess could lead to similar phenotypes such as stem height stimulation (**Figure 2C**), hypocotyl xylem II to total xylem ratio (**Figure 3B, C**) and others observed in this study. Moreover, the upregulation of *ERECTA* and *BAK1* in hypocotyls (**Figure 8D**) could provide direct explanation for the observed reduction in xylem II (**Figure 3 B,C**). Although the signaling cascades involving these different receptor like kinases are currently not known, clearly the cell wall integrity signaling could be responsible for many XET effects that are indirect and non-cell autonomous as well as those involving microtubule network modulation.

### XET as a target for biomass improvement

Considering the extensive effects of XET on secondary growth revealed by this study, XETs may become targets for woody biomass improvement. In support, our analysis demonstrated 15% gains compared to wild type in sugar production in *xth9* mutants (**Figure 5 F**), which also exhibited over 50 % growth increase (**Figure 2 C**). Our data with Arabidopsis XET-modified genotypes indicate that several wood traits are affected by XET. Indeed, Wang and Zhang (2014) identified single nucleotide polymorphisms in *PtoXTH34* in *P. tomentosa* associated with lignin content, stem volume, and MFA, which explained up to 11.0% of the phenotypic variance. Our data in Arabidopsis suggest causality behind these associations.

The results with XET affected plants indicate that other manipulation of XG metabolism could lead to strong effects on wood properties. This conclusion is supported by studies with the constitutive overexpression of *Aspergillus aculeatus XYLOGLUCANASE 2* (*AaXEG2*) in trees, which ectopically decreases XG content (Park et al., 2004). Such modified trees exhibited several developmental alterations, including stem height and diameter increase (Park et al., 2004; Hartati et al., 2011), increased cellulose content, density and elastic modulus (Park et al, 2004), increased cellulose microfibril width (Yamamoto et al., 2011), and remarkable improvement in saccharification for both glucose yields (50% gains) and cellulose conversion (60% gains) (Kaida et al., 2009). Such gains are among the highest reported so far for hemicellulose-modified tree species (Donev et al., 2018). Thus, the manipulation of XG metabolism is a vital route for tree improvement despite of its low content in wood cell walls.

## METHODS

### Plant materials and growth conditions

Sterile seeds of *Arabidopsis thaliana* Col0 (wild-type), transgenic plants and the mutants were vernalized in 0.1% agar at 4°C for three days and planted in soil with vermiculite (3:1). Plants were grown at 22°C and light intensity 120-150 µEm^-2^s^-1^ under long day conditions (16 h light /8 h light) for six-eight weeks. For phenotyping secondarily thickened hypocotyls, plants were first subjected to short days (8 h light /16 h dark) for four weeks and then to long days for additional two-four weeks.

### Production and selection of T-DNA insertion lines and *PtxtXTH34* OE lines

T-DNA mutant insertion lines (SAIL 681-G09 and SAIL 722-B10) (**Figure 2A**) obtained from NASC were back-crossed several times until the single locus (as determined by the segregation ratio), homozygotic plants were obtained. OE lines carrying *PtxtXTH34* were obtained as described previously (Miedes et al., 2013).

### Histochemical β-glucuronidase activity tests for promotor activity studies

pXTH4::GUS plants were obtained by transforming Col0 with the vector containing 5′ upstream sequence (1 kb) from the start codon of *XTH4* (At2g06850). The primers used were: 5′-cacc TAT TTT GAT AGT AGA GAG TTG GAA TAC CAT-3′ and 5′-AAC AGT CAT GGT TGT GTT TTA GAA AGA GAT-3′. The fragment was inserted into pBGGUS (Kubo et al., 2005). pXTH9::GUS line was selected by Becnel et al., (2006). At least three independent lines were analyzed before selecting the representative line for final analyses. Cross sections of the mature hypocotyls and basal parts of the stems of pXTH4::GUS and pXTH9::GUS lines were incubated for 4 h in a mixture of 1 mM X-Gluc, 1 mM K₃3Fe(CN)₆, 1 mM K₄4Fe(CN)₆, 50 mM sodium phosphate buffer pH=7.2, and 0.1% Triton X-100 at 37°C, fixed in FAA (5% formaldehyde, 5% acetic acid and 50% ethanol) for 10 min, rinsed several times in 50%, and then in 100% ethanol until the chlorophyll was completely removed, and observed under an Axioplan 2 microscope connected with an AxioVision camera (Carl Zeiss, Germany).

### XET-activity by radiometric tritium assay

Stems and hypocotyls of three 6-week-old plants were ground in dry ice and then added to 0.5 M NaCl (0.5 mg tissue/1 µL salt). The slurry was centrifuged at 5000xg for 5 minutes at 4°C and supernatants were analyzed for XET activity using radiometric assay (Fry et al., 1992). 10 µL of each extract were incubated for 2 h at room temperature in a volume of 40 µL of reaction mixture containing 50 mM MES-Na, pH 6.0, 1.4 kBq tritium labelled XLLGol, and 2 mg/ml XG. The reaction was stopped by adding 100 µL of 20% formic acid. The resulting 140 µL mixture was spotted onto 5 cm x 5 cm piece of 3M paper and allowed to air dry overnight. The 3M paper was washed with running water at ∼45 degree angle for 2 h and air dried. The scintillation was carried out with 2 ml scintillant in a Tri-Carb 2100 liquid scintillation analyzer. Five biological replicates (pools of three plants) were used for each line.

### Immunodetection of XG

The basal parts of the inflorescence stems and hypocotyls of 6-week-old plants were cross-sectioned at 60 µm thickness using a vibratome (VT 1000S, Leica, Germany), and fixed in 4% paraformaldehyde dissolved in 0.5M PIPES buffer, pH 7.0 for 30 min. The sections were then washed 5 times for 3 min in PBS with 0.1% Tween 20 (PBS-T), blocked in 5% skim milk in PBS buffer for 1 h at room temperature, incubated in primary antibody CCRC-M1 (gift from Dr. Michael Hahn) for 2 h at room temperature, washed 5 times for 3 min in PBS-T and incubated in secondary antibody Alexa Fluor 488 anti-mouse gold for 2 h in the dark. After removal of the secondary antibody by washing 5 times for 3 min in PBS-T, the sections were observed by a laser scanning confocal microscope (LSCM) Zeiss LSM 510 (Jena, Germany) with a 488-nm argon-krypton laser.

### Histological analysis

The hand cross-sections of mature hypocotyls were stained with the 1% Phloroglucinol in 18% HCl, and viewed under the Axioplan 2 microscope equipped with an AxioVision camera (Carl Zeiss, Germany). The proportions between tissues were measured for 10 hypocotyls from each line using image analysis (Axio Vision, Sweden). Four measurements per section were made along four equidistant radii around the hypocotyl circumference.

### Wood maceration

10 hypocotyls of each line were put in a tube and macerated in a mixture of 3% H_2_O_2_ and 50% glacial acetate acid at 95 °C for 8 h, washed in water and neutralized. The individual cells were released by vigorous shaking of tubes and viewed with the differential interference contrast in an Axioplan microscope (Carl Zeiss, Germany). For each analyzed line, the length and width of 100 fibers and 300 vessel elements were measured using Axio Vision image analysis software (Carl Zeiss, Germany).

### Transmission Electron Microscopy (TEM)

Approx. 3 mm long segments from basal part of the inflorescence stems and from hypocotyls of two 8-week-old plants per each line (WT, *xth4*, *xth9*, *xth4x9* and OE18) were fixed in 4% paraformaldehyde and 0.05% glutaraldehyde (Analytical standards AB, Mölnlycke, Sweden) in 25 mM phosphate buffer, pH 7.2. They were dehydrated in a graded ethanol series and embedded in LR-White resin (TAAB, Berkshire, UK) and polymerized at 60°C for 16-20 h. Cross sections were cut using ultramicrotome and observed with a JEOL JEM-1230 transmission electron microscope (JEOL, Tokyo, Japan). Measurements of the cell wall thickness (without the compound middle lamella) were performed in 100 cells (50 cells per plant) for hypocotyl fibers and 15 cells (7-8 cells per plant) for the other cell types, at all contacts with neighboring cells, and counts of secondary cell wall layers were performed for 50 cells of each type (25 cells per plant), including interfascicular fibers (IF), fascicular fibers (FF), fascicular vessel elements in the stems, and secondary xylem fibers (HF) and vessel elements in hypocotyls.

### Cell wall analyses

Approx. 30 d old plants having 20 cm long stems were used for cell wall analyses. Basal 10 cm long segments were collected and pooled to make one biological replicate. They were immediately frozen in liquid N_2_, freeze-dried for 36 h and ball milled to fine powder in stainless steel jars with one 12 mm grinding ball at 30 Hz for 2 min, using an MM400 bead mill (Retsch). The alcohol insoluble residue (AIR1) was prepared from wood powder as described earlier and analyzed by pyrolysis GC/MS and FT-IR spectroscopy (Gandla et al., 2015). AIR1 material was destarched to obrain AIR2, and used for determination of monosugar composition after methanolysis and trimethylsilyl/TMS derivatization, and cellulose content (Gandla et al., 2015). AIR1 and AIR2 fractions were used to analyze lignin content (Foster et al., 2010).

### Vibrational microspectroscopy

0.5 cm segments from hypocotyls and from basal part of side branches from six-week-old plants were immediately frozen in liquid N_2_ and stored in −20 °C. Frozen material was mounted in O.C.T. compound (361603E VWR Chemicals) and sectioned using HM505E (Microm) cryostat. 10 µm thick cross-sections were washed in water and transferred to a Raman and infrared transparent CaF_2_ slides. The sections were dried in a desiccator for at least 48 hours. The same section was analyzed by *in situ* FTIR and Raman microspectroscopy.

#### FTIR microspectroscopy

FTIR microspectroscopic measurements and data analysees were performed as described by Gorzsás et al. (2011). Spectra were recorded using 32 scans at a spectral resolution of 4 cm^−1^ on a Bruker Tensor 27 spectrometer equipped with a Hyperion 3000 microscopy accessory, including a liquid nitrogen cooled 64 × 64 mercury cadmium telluride focal plane array detector (Bruker Optik GmbH). Five plants were used for each line, and 4-7 positions were selected in each section for hyperspectral imaging, representing entire cross sections. Suboptimal quality images (such as saturated spectra due to folding upon drying, etc.) were excluded from further analyses. Ten spectra / image and cell type were extracted, compiled and exported as .mat files using OPUS (version 7.0.129, Bruker Optik GmbH). The spectral region of 900–1800 cm^−1^ was used in subsequent analyses.

#### Raman microspectroscopy

Raman microspectroscopy (mapping) was performed as described by Gorzsás (2017) on a Renishaw InVia Raman spectrometer equipped with a CCD detector, using a 514 nm Ar^+^ ion laser with 2s irradiation time and 100 percent laser power set by the software (WiRE, versions 3.0-3.4, Renishaw Plc, UK), in static mode, centered at 1190 cm^-1^ (resulting in a spectral region of ca. 510 - 1803 cm^-1^ with ca. 1 cm^-1^ spectral resolution using a 2400 lines grating). A 50x magnification lens was used, with 1.5 micrometer step sizes (XY direction), in normal confocal mode, recording images consisting of a minimum of 100 spectra / image. Raman shifts were calibrated using the built-in Si standard of the instrument. Cosmic rays were removed and the hyperspectral data noise filtered using the chemometrics package of WiRE (version 3.4, Renishaw Plc, UK). Five plants were analysed for each line, recording two independent hyperspectral images / plant, and using five (FF and HF) or 10 (IF) spectra / image for multivariate analyses. Suboptimal quality images were excluded.

#### Processing and data evaluation of vibrational spectra

Spectra were pre-processed using the open-source MCR-ALS script (run in Matlab environments, versions 14a-18a, Mathworks Inc, USA), as provided by the Vibrational Spectroscopy Core Facility (http://www.kbc.umu.se/english/visp/download-visp/). Baseline correction was performed using asymmetrical least squares (Eilers, 2004), with λ varying between 100,000 (Raman spectra) and 10,000,000 (diffuse reflectance FTIR spectra), while p was always kept at minimum (0.001). Spectra were smoothed by Savitzky-Golay filtering (Savitzky and Golay, 1964), using a first order polynomial and a frame rate of five. Spectra were total area normalized in the spectral region used for the analyses and analyzed by SIMCA-P (versions 13.0-14.0, Umetrics AB Sweden). Spectral data were centered, while Y variables (average cell wall layer numbers) were unit variance scaled. Initial PCA models were created using the “Autofit” function for overview purposes and identifying potential outliers before OPLS-DA models. OPLS-DA models used a fixed number of components, as listed for each model, kept to a minimum to avoid overfitting.

### Gene expression analyses

The total RNA was extracted from frozen ground basal stem and hypocotyl tissues using RNeasy^®^ Plant Mini Kit (Qiagen), DNAse treated using DNA-free™ Kit (Invitrogen) and cDNA was synthesized using iScript™ cDNA synthesis kit (BIO-RAD) following the manufacturer’s protocol. The quantitative Real Time-PCR (qRT-PCR) was performed using Light Cycler^®^ 480 SYBR® Green I master mix (Roche) in a Light Cycler 480 II instrument (Roche) as per the manufacturer’s instructions. Primers used for qRT-PCR are listed in **Table S1**.

## ACKNOWLEDGEMENTS

We thank Dr. Takahisa Hayashi for helpful comments on the manuscript, Dr. Michael Hahn for CCRC-M1 antibody and Dr. Hans Stenlund (SLU) for advice on analyses of microspectroscopic data, Nottingham Arabidopsis Seed Centre for the SAIL lines, and the Umeå Core Facility for Electron Microscopy (UCEM) for help and advice.

## SUPPLEMENTAL DATA

**Supplemental Figure 1.** Identification of secondary growth-related XTH genes in *P. trichocarpa* and in *A. thaliana*.

**Supplemental Figure 2.** Effect of altered XET level on cell wall ultrastructure in inflorescence stems fascicular and interfascicular fibers as visualized by Transmission Electron Microscopy (TEM).

**Supplemental Figure 3.** Effect of altered XET level on cell wall ultrastructure in inflorescence stem xylem vessels elements as visualized by Transmission Electron Microscopy (TEM).

**Supplemental Figure 4.** Effect of altered XET level on cell wall morphology in hypocotyl xylem vessel elements as visualized by Transmission Electron Microscopy (TEM).

**Supplemental Figure 5.** OPLS-DA models (pairwise comparisons) based on diffuse reflectance FTIR spectra from basal inflorescence stem alcohol insoluble residue of different genotypes.

**Supplemental Figure 6.** I*n situ* FTIR microspectroscopy of cell walls in fibers of inflorescence stems and hypocotyls in *xth* mutants and overexpressing plants.

**Supplemental Figure 7.** OPLS-DA models (pairwise comparisons) for interfascicular fibers based on Raman microspectroscopic data.

**Supplemental Table 1.** List of primers used in qPCR.

